# *bsAS*, an antisense long non-coding RNA, controls cell fate through regulation of *blistered/DSRF* isoform expression

**DOI:** 10.1101/846451

**Authors:** Sílvia Pérez-Lluch, Alessandra Breschi, Cecilia C. Klein, Marina Ruiz-Romero, Amaya Abad, Emilio Palumbo, Lyazzat Bekish, Carme Arnan, Roderic Guigó

## Abstract

**Summary:** Natural Antisense Transcripts (NATs) are long non-coding RNAs (lncRNAs) that overlap coding genes in the opposite strand. NATs roles have been related to gene regulation through different mechanisms, including post-transcriptional RNA processing. With the aim to identify NATs with potential regulatory function during fly development, we generated RNA-Seq data in eye-antenna, leg, and wing at third instar larvae. Among the candidate NATs, we found *bsAS*, antisense to *bs/DSRF*, a gene involved in wing development and neural processes. Through the analysis of the RNA-Seq data, we found that these two different functions are carried out by the two different protein isoforms encoded in the *bs* gene. We also found that the usage of these isoforms is regulated by *bsAS*. This regulation is essential for the correct determination of cell fate during *Drosophila* development, as *bsAS* knockouts show highly aberrant phenotypes. *bs* regulation by *bsAS* is mediated by the specific physical interaction of the *bsAS* promoter with the promoters of *bs*, and it likely involves a mechanism, where expression of *bsAS* leads to the collision of RNA polymerases acting in opposite directions, preventing the elongation of the longer isoforms of *bs*, the ones carrying the neural related functions. Evolutionary analysis suggests that the *bsAS* NAT emerged simultaneously to the long-short isoform structure of *bs*, preceding the emergence of wings in insects, and maybe related to regulation of neural differentiation.

## Main text

The *D. melanogaster* genome encodes 16,698 genes, including 13,920 protein coding genes, 2,433 lncRNAs and 308 pseudogenes (FlyBase v6.05 (Attrill et al., 2016)). Although antisense lncRNAs are not explicitly annotated, they can be inferred from their overlap and orientation relative to protein coding genes. We calculated that 855 lncRNAs (35%) overlap 873 protein coding genes in antisense orientation (Natural Antisense Transcripts, NATs), forming 953 sense-antisense pairs, a number in line with previous reports (Lapidot and Pilpel, 2006; Sun et al., 2006). The coding (sense) and non-coding (antisense) gene pairs (SA pairs), can be arranged in different configurations (Figure 1A): in most of the pairs, the lncRNA is fully included within the protein coding gene (391 pairs, 41%), followed by 5’ head-to-head pairs (165 pairs, 17%) and 3’ tail-to-tail pairs (61 pairs, 7%). Only in a minority of the pairs (28 pairs, 3%), the protein coding gene is fully included in the lncRNA, while the remaining 32% of the pairs (308) are arranged in complex configurations involving three or more genes (Figure 1A). Remarkably, fly protein coding genes involved in SA pairs are significantly associated to developmental and morphogenetic processes (Figure 1B). This strongly suggests that NATs might play a relevant regulatory role in fly development.

**Figure 1.**
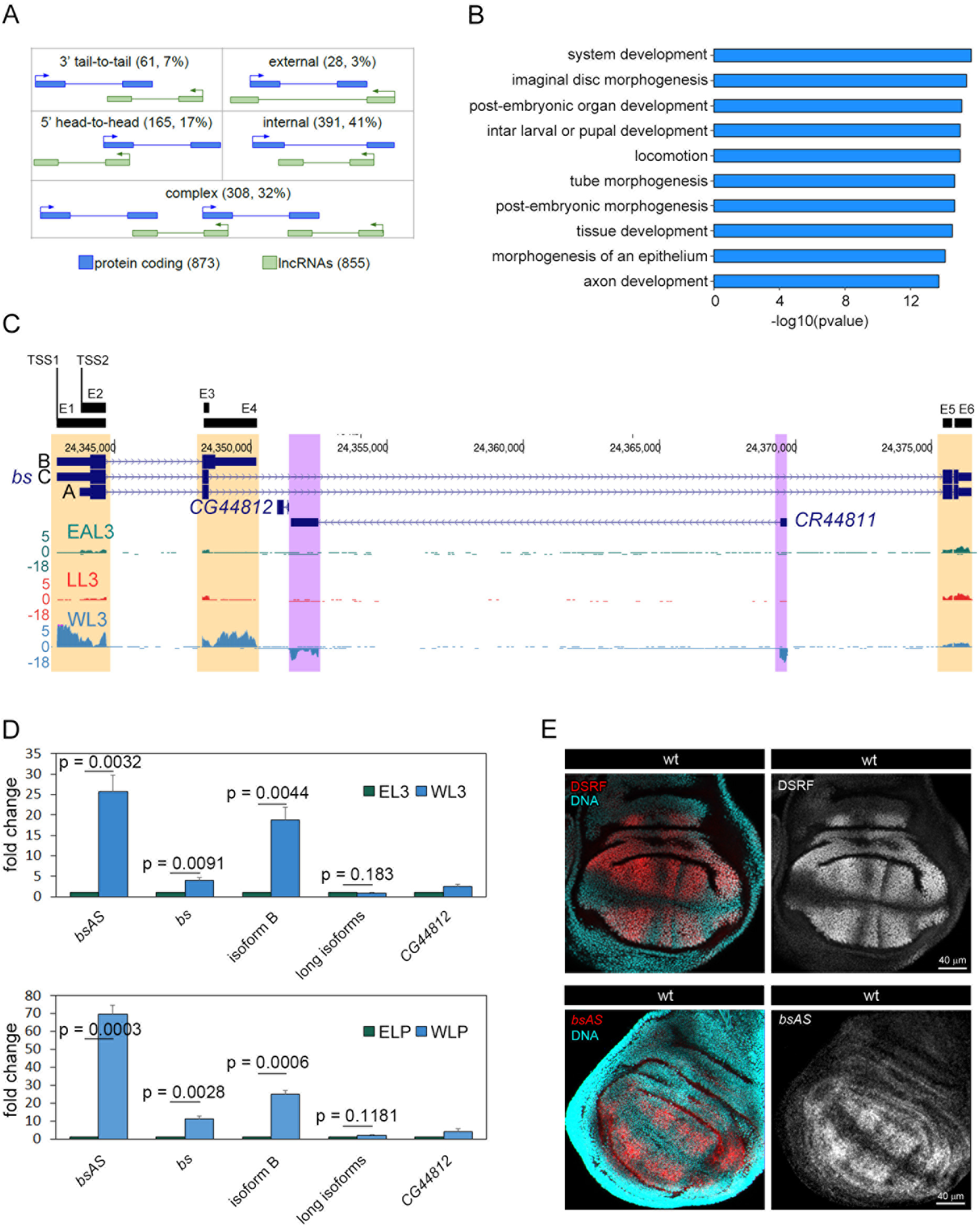
Natural Antisense Transcription in *Drosophila melanogaster*. See also Supplementary Figure 1 and Supplementary Tables 1 and 2. **A.** Configuration of sense/antisense pairs identified in *D. melanogaster* genome. We have identified 855 lncRNAs overlapping 873 coding genes in the fruit fly annotation. Around a 32% of the SA pairs present a complex configuration, meaning that one lncRNAs overlaps more than one coding gene or the other way around. **B.** Gene Ontology Term Enrichment analysis of coding genes overlapping antisense transcripts. Sense genes are mainly related to development and morphogenesis categories. **C.** Expression of the *bs* locus in the different tissues. Isoform B of *bs* is expressed mainly in wings, correlating with the highest expression of the AS lncRNA *CR44811*. In the other tissues, the isoform A seems to be the predominant one. In the leg, also some expression of the lncRNA is observed. Regions showing differences in expression between tissues are highlighted (*bs* TSSs in green, *bs* Transcription Termination Sites –TTSs– in blue and AS in purple). **D,** Relative expression of *bs* and *bsAS* in wings and eyes at third instar larvae (L3 –upper panel–) and late pupa (LP –lower panel–) stages. *bsAS* and the short isoform B of the coding gene are much higher expressed in wings than in eyes, whereas there are not significant differences in the expression of the long isoforms. The protein coding gene *CG44812*, of unknown function, is almost not expressed in these tissues. Error bars depict Standard Error of the Mean (SEM) of at least three biological replicates. Statistical significance was computed by one-sample t-test. **E,** Expression pattern of DSRF and *bsAS* in wt wing imaginal discs. Upper panels, immunostaining of DSRF (red and grey) in wt WL3. DSRF is expressed in the intervein region of the wing pouch. Lower panels, *in situ* hybridization of *bsAS* (red and grey) in WL3. The expression of *bsAS* mimics DSRF expression pattern in the intervein regions. Nuclei, stained with DAPI, are depicted in blue in all cases.

With the aim to identify NATs with potential regulatory function during fly development, we performed strand-specific RNA-Seq in biological replicates on three imaginal tissues of *D. melanogaster* third instar larvae (L3): eye-antenna (EAL3), leg (LL3) and wing (WL3) (Supplementary Figure 1A and Supplementary Table 1). Of the 953 SA pairs, we found 145 (about 15%) in which the two members of the pair were detected with more than 0.5 TPMs (Transcripts per Million Mapped Reads in at least one tissue, albeit not necessarily the same). Since splicing regulatory activity has been documented for a human NAT (Beltran et al., 2008) and extensive relationship between antisense transcription and splicing has been hypothesized in human (Morrissy et al., 2011), we explored specifically the relationship between NAT expression and alternative transcript usage across fly larval samples. Of the 145 SA pairs with both genes expressed in at least one tissue in L3, 104 pairs (72%, 102 genes) involve a protein coding gene with multiple isoforms. We compared the expression of these NATs with the inclusion of the 964 exons of the sense protein coding genes, focusing on the exons the inclusion of which changed the most (see Methods). We found a few cases of relationship between expression of the NAT and the splicing of the sense protein coding genes—the most notable being the case of *blistered* –*bs* (Supplementary Figure 1B).

In *Drosophila melanogaster*, the *blistered* gene (*bs*) gene is essential, among other functions, for the proper development of wings (Roch et al., 1998), and it is also involved in neural processes (Donlea et al., 2009; Thran et al., 2013). The protein encoded by *bs*, *Drosophila* Serum Response Factor (DSRF), is present in two different isoforms in the fly. They both carry the DNA binding domain MADS-box, but differ at the terminal end, one much longer than the other (449 vs. 355 amino acids). The long isoform is encoded by two transcripts (variants A and C in Figure 1C), that use two different transcription start sites (TSS2 and TSS1, respectively), and differ only in the first exon. The other protein isoform is encoded by a shorter two-exon transcript (variant B), which shares TSS1 with isoform C. The two long isoforms contain a 26 Kb intron that encompasses the NAT, *CR44811* (herein *blistered AntiSense* – *bsAS*) and a coding gene of unknown function (*CG44812*).

There is a contrasting pattern of *bs* isoform expression between wing and eye and leg tissues (Figure 1C). In wing, we observe the predominant expression of short isoform B, while in eye-antenna and in leg we observe exclusive expression of the long isoform A. We also detect some expression of the long isoforms in wing imaginal discs (which we hypothesize is originating from isoform A in the vein regions). The NAT *bsAS* also shows contrasting expression, being very highly expressed in wing—where is one the most highly expressed lncRNAs in the fly genome (Supplementary Figure 1C)—but very low in eye-antenna and leg imaginal discs. We independently confirmed the expression of the *bs* isoforms and of *bsAS* by qPCR in wing and eye at third instar larvae and late pupa stages (Figure 1D). The restricted expression of *bsAS* to wing imaginal discs is also supported by the presence of trimethylation of lysine 4 of histone H3 (H3K4me3, mark associated to active promoters) in wing, but not in eye imaginal discs (Supplementary Figure 1D-E). In situ hybridization in wing imaginal discs reveals that *bsAS* exhibits the same spatial expression pattern as *bs*, restricted to the intervein regions (Figure 1E). In summary, thus, in wing intervein regions (non-neural tissues), *bsAS* is highly expressed, and TSS1 is actively driving the expression of the short isoform B. In contrast, in eye and, possibly, wing veins (neural tissues), *bsAS* is not expressed, and TSS2 is actively driving the expression of the long isoform A.

All these observations suggest that *bsAS* could play a role in regulating isoform usage of *bs*. To test this hypothesis, we used CRISPR/CAS9 to induce a 560 bp deletion surrounding the TSS of *bsAS* (Figure 2A and Supplementary Figure 2A-C). This effectively abolishes expression of this gene in L3 and LP in wing imaginal discs (Figure 2B). Knocking out *bsAS* has little effect on the expression of the short isoform B, but induces overexpression of one (or both) of the long isoforms. As a result, there is an overall increase in the expression of *bs* in the *bsAS* mutant wings compared to wt (particularly at protein level, compare Figure 2C, to Figure 1E upper panel). Deletion of the *bsAS* promoter did not alter the pattern of expression of DSRF, restricted to the intervein regions of the wing disc (Figure 2C). Animals carrying the deletion, however, exhibited notable defects in wing development, much stronger in homozygous mutant flies, and resembling those of the *bs* mutant flies (Fristrom et al., 1994; Montagne et al., 1996; Roch et al., 1998) (Figures 2D-F). These were blistered, presenting necrotic regions and strong defects in vein/intervein patterning, in particular, extra vein tissue in intervein regions. Ectopic expression of *bsAS* did not rescue the wing phenotype in the mutant (Figure 2G and Supplementary Figure 2D), suggesting a role for *bsAS in cis*, likely due to the transcription process rather than to the *bsAS* transcript itself. The phenotype is a consequence of the increase in expression of the long isoforms of *bs* in the *bsAS* mutant, since similar wing defects are observed when overexpressing the DSRF long isoform in the wing pouch (Figure 2H and Supplementary Figures 2E-F). Overexpression of the short isoform B, on the other hand, induces strong defects in wing development and even animal death (Figure 2I and Supplementary Figures 2G-H), suggesting that the short DSRF protein is toxic when expressed in excess or in tissues where it is not canonically expressed.

**Figure 2.**
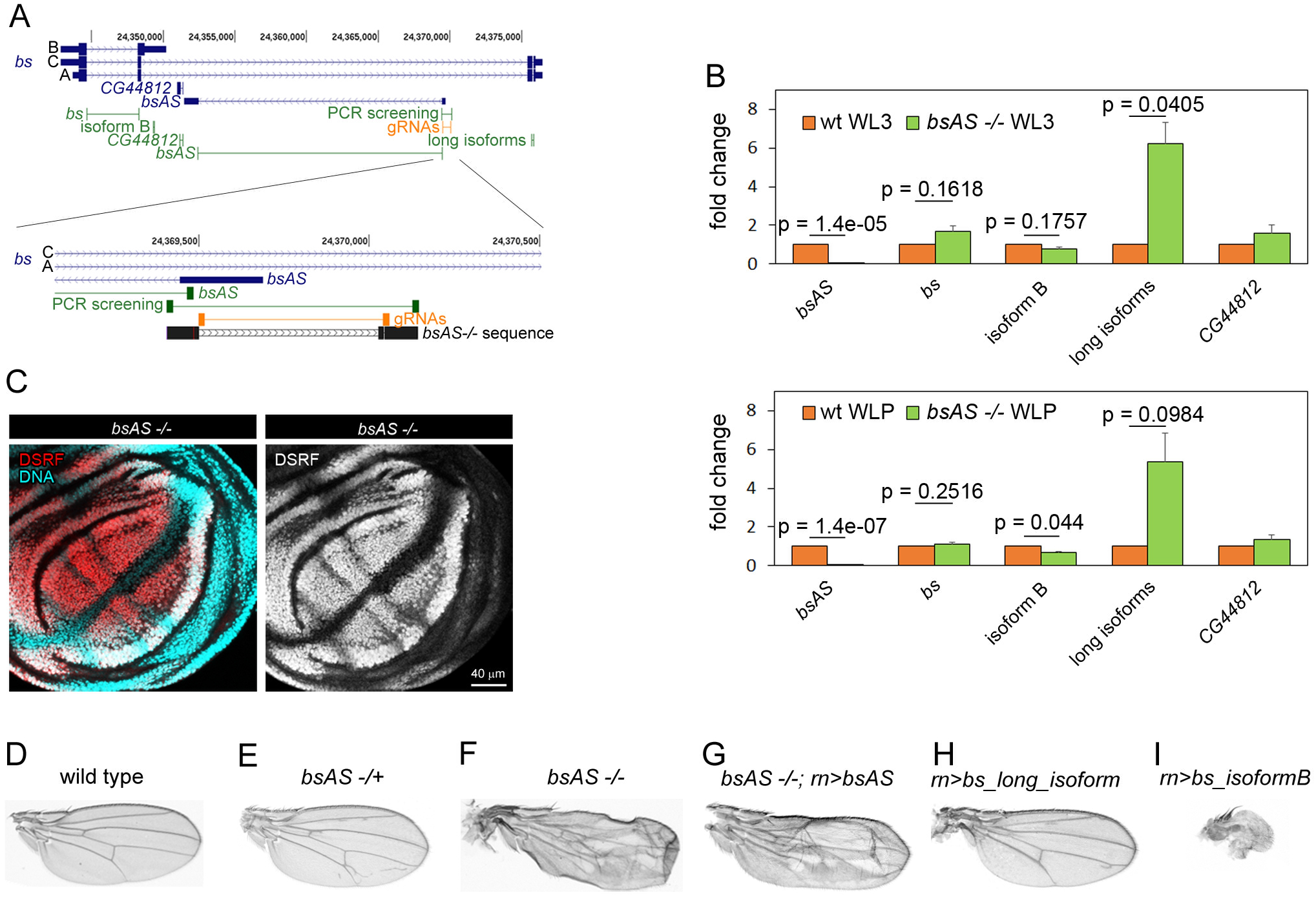
*bsAS* controls *bs* isoform usage during fly development. See also Supplementary Figure 2 and Supplementary Tables 1, 2 and 3. **A,** Distribution of primers and CRISPR gRNAs along the *bs* locus. Upper panel, primers amplifying the different isoforms of *bs* and *bsAS* are depicted in green; gRNAs used to induce the deletion of *bsAS* are depicted in orange. Lower panel, zoom of the lncRNA TSS region. Sanger sequence of *bsAS* mutant flies is depicted in black. A long 560 bp deletion was induced around the TSS of *bsAS*. A small deletion of 2 nucleotides at the 5’ region of the TSS is also observed. **B,** Relative expression of *bs* and *bsAS* in *bsAS* −/− and wt L3 (upper panel) and LP (lower panel) wings. The lncRNA is not expressed in *bsAS*−/− wings. The overall expression of the *bs* gene seems to be slightly higher in mutant than in wt wings, especially in L3 stage; however, when comparing the expression of *bs* isoforms, we do not observe differences in the expression of the short one between the mutant and the wt, while the long isoforms are overexpressed 6-fold in the mutants. Error bars depict Standard Error of the Mean (SEM) of at least three biological replicates. Statistical significance was computed by one-sample t-test. **C,** Expression pattern of DSRF *bsAS*−/− in wing imaginal discs. Immunostaining of DSRF (red and grey) in *bsAS*−/− WL3. No differences in DSRF expression pattern are observed between wt and *bsAS-/-* L3 wings (Fig. 1c). Nuclei, stained with DAPI, are depicted in blue in all cases. **D-I,** Adult wings from males. **D,** Wild type. **E,** *bsAS* heterozygous mutant. The wings present extra veins invading intervein tissue. **F,** *bsAS* homozygous mutant. The wings are creased and vein pattern is completely impaired. **G,** *bsAS* homozygous mutant overexpressing transgenic *bsAS* under the control of *rotund* (*rn*) driver. The mutant phenotype is not rescued by the overexpression of the lncRNA. **H,** Wings overexpressing *bs* isoform A under the control of the *rn* driver. Extra veins appear within intervein tissue, resembling the phenotype of *bsAS* heterozygous mutants. **I,** Wings overexpressing *bs* isoform B under the control of the *rn* driver. This overexpression induces lethality and strong abnormalities in adult wings.

To monitor the molecular changes underlying the *bsAS* mutant phenotype, we performed RNA-Seq of mutant wing imaginal discs in L3 and LP (Supplementary Figure 2I and Supplementary Table 2). Results indicate that the long isoform specifically induced in the *bsAS* mutant is isoform C, while the expression of isoform A is not affected. We observed few genes differentially expressed comparing the mutant with the wt flies in L3 (27 up and 29 down regulated), but many more in LP: 275 upregulated and 260 downregulated genes (Supplementary Table 3). There were not clear functional categories enriched among downregulated genes (Supplementary Figure 2J) but, after closer inspection, we identified many genes involved in wing morphogenesis and cell adhesion among them. These include *Sox102F*, that regulates the expression of the *Wnt* pathway and has been related to wing vein development and patterning (Li et al., 2013), *ImpE2* and *ImpE3* that are ecdysone-inducible genes related to imaginal disc eversion (Andres et al., 1993; Moore et al., 1990; Paine-Saunders et al., 1990) and *multiple edematous wings* (*mew*) and *miniature* (*m*), that are genes related to cell adhesion and wing morphogenesis (Brabant et al., 1996; Roch et al., 2003). Upregulated genes were strongly enriched in functions related to photo transduction, dopamine biosynthesis and other neural specific functions (Supplementary Figure 2K). Among the genes upregulated in mutant wings we can find *Dscam1* and *Dscam4*, cell adhesion molecules related to axon guidance and neural development (Schmucker et al., 2000; Tadros et al., 2016), and *inaC*, *inaD* and *ninaE*, genes involved in the detection and response to light stimuli (for a review see (Pak et al., 2012)).

All together these results suggest that the expression of the *bs* long isoforms induces the expression of neural specific genes, and that *bsAS* prevents tissue neuralization in the wing intervein regions by repressing specifically the long isoform C, and promoting, in consequence, the expression of short isoform B. Since these two isoforms share TSS1, we investigated whether physical interactions between the TSS of *bsAS* and TSS1 could underlay *bsAS* regulation of *bs* isoform usage. Using publicly available HiC data in Kc167 cells (Cubenas-Potts et al., 2017), we did find an enrichment in contacts between the TSS of *bsAS* and TSS1, even though *bs* and *bsAS* are poorly expressed in these cells (Duff et al., 2015) (Figure 3A and Supplementary Figure 3A). We further performed 3C assays in L3 wing and eye imaginal discs, interrogating the interaction of *bsAS* TSS with 10 different regions along the *bs* locus (including TSS1 and TSS2) (Figure 3B). We detected significant interactions with TSS1 both in wing and eye, but not with TSS2 (Figure 3C). The interaction seems to be very stable thus, present even in tissues in which TSS1 and *bsAS* are inactive. We detected a second interaction with a region known to be a hot spot for transcription factor binding in fly embryos (Washington et al., 2011) (Supplementary Figure 3B). Interactions between TSS1 and a region adjacent to *bsAS* TSS, which were maintained in wt wings, were dramatically impaired however in the *bsAS* mutant (Figure 3D), indicating that sequences within this region are likely responsible for the interaction with TSS1. The only factor for which binding sites were found both at the TSS1 and *bsAS* TSS was the GAGA factor (GAF), for which no binding sites were found at TSS2. GAF ChIP-Seq data available for L3 wing imaginal discs (Oh et al., 2013) revealed strong GAF peaks at TSS1 and *bsAS* TSS (Supplementary Figure 3C), providing additional support for a role of GAF in mediating the interaction. Indeed, the downregulation of the *Trithorax-like* (*Trl*) gene, encoding for GAF, induces a reduction of the intervein regions (Blanch et al., 2015) and the appearance of extra vein tissue in adult wings (Bejarano and Busturia, 2004), confirming the role of GAF in vein/intervein patterning during wing development. The loop formed by this interaction is likely stabilized by cohesin complexes, which have been demonstrated to contribute to interactions between chromatin regions such as enhancer-promoter communication (Dorsett, 2019). We also observed two strong Pol II peaks (Schertel et al., 2015) at both TSSs in L3 wings, confirming that the two sites are transcriptionally active in this tissue at this developmental stage.

**Figure 3.**
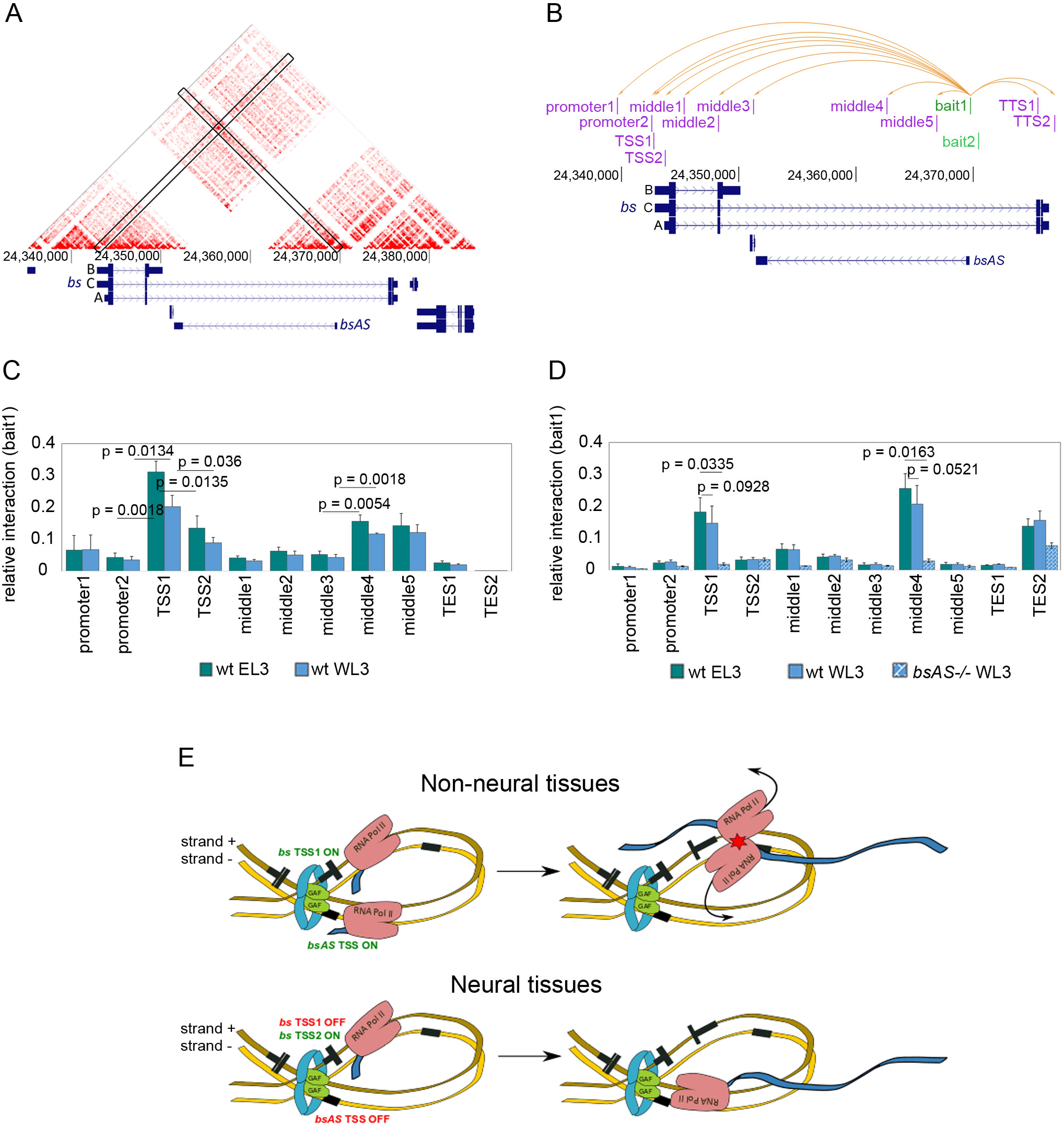
*bsAS* and *bs* are co-regulated through the interaction of their TSSs. See also Supplementary Figure 3. **A,** JuiceBox tool (Durand et al., 2016) was used to visualize high-resolution HiC maps generated in Kc167 cells (Cubenas-Potts et al., 2017). A strong interaction signal is observed in the intersection between *bs* TSS1 and *bsAS* TSS. **B,** Representation of *bs* locus and the tested interactions inside this region. Two different baits were anchored at *bsAS* TSS (in green) and interaction enrichment was assessed for each bait by 3C experiments against all depicted regions (in pink). Orange arrows represent all tested interactions for bait1. The same interactions were tested for bait2. **C,** Interaction between bait1 and *bs* locus in wild type wings and eyes. Interaction enrichment is significantly higher between *bsAS* TSS and TSS1 than between the *bsAS* TSS and the other tested regions, both in wings and eyes. A significant increase in relative interaction is also detected between *bsAS* TSS and middle regions 4 and 5. **D,** Interaction between bait2 and *bs* locus in wild type wings and eyes and *bsAS*−/− wings. The interaction between bait2 and TSS1 is still enriched in wild type tissues, but it is heavily impaired in *bsAS* mutant wings. Also the interaction between *bsAS* TSS and middle4 region is lost in the mutant background. In **C** and **D**, error bars represent the SEM of at least three biological replicates. Statistical significance was computed by two-tailed t-test. **E,** In both panels, positive and negative DNA strands are depicted as dark and light brown ribbons, respectively. Transcribed regions are depicted in black, narrow bars representing non-coding regions and wide bars representing coding regions. For *bs*, only the active TSS is depicted in each case. Blue ribbons represent nascent RNA and blue rings represent cohesion proteins. *bsAS* TSS and *bs* TSS1 interact in a highly stable manner, independently of the expression of the coding and the non-coding gene. Upper panel. In non-neural tissues, *bs* TSS1 and *bsAS* TSS are active and RNA Polymerases are recruited in both strands. Elongation along the two strands provokes the eventual collision of the polymerases, transcription is impaired and only the short isoform of *bs* can be fully transcribed. Lower panel. In neural tissues, such as eye-antenna imaginal disc or the vein regions of the wing disc, *bs* TSS1 and *bsAS* TSS are inactive; only *bs* TSS2 is active. RNA Polymerase II is recruited only in the positive strand allowing for the polymerase to elongate until the end of the locus, and the long isoform can be fully transcribed.

All these observations suggest a model to explain how *bsAS* regulates the isoform usage of the *bs* gene and represses neural differentiation (Figure 3E). In non-neural tissues, *bs* TSS1 and *bsAS* TSS are active, and RNA Polymerases are recruited in both strands. TSS1 triggers the transcription of both the short isoform B and the long isoform C, but the elongation along both strands provokes the eventual collision of the polymerases and, in consequence, long isoform C cannot be generated. Only the short DSRF protein is generated, and neural differentiation does not occur. In neural tissues, only TSS2 is active, the RNA polymerase is only recruited at the sense strand, and the elongation of the long isoform A is not compromised. The long DSRF protein is generated, leading to neural differentiation.

We did not find any known amino acid motif in the 111-aa long region specific to the DSRF long isoform, that could explain its specific neural function. Actually, outside of the MADS box, which is strongly conserved across metazoans (and, intriguingly, systematically interrupted by an intron, in spite of dramatic changes in the exonic structure of the *bs* gene through evolution), there is little conservation of the DSRF protein, even already within diptera (Figure 4 and Supplementary Figure 4). Evolutionary analysis, however, showed that the long-short isoform structure of the *bs* gene appeared much earlier, at the root of pancrustacea (hexapoda and crustacea, Figure 4A). By using available RNAseq data, we have been able to trace the origin of *bsAS* also at the root of pancrustacea (Figure 4B and Supplementary Figure 4B), suggesting that both the long-short isoform structure of *bs* and *bsAS* emerged simultaneously. Since crustaceans and non-insect hexapods are wingless animals, *bsAS* regulation of *bs* isoform usage precedes the emergence of wings in insects, and it may be primarily related to neural differentiation, since, as in the fly, we have observed through insects low *bsAS* expression in the brain compared to whole body expression (Figure 4C-D).

**Figure 4.**
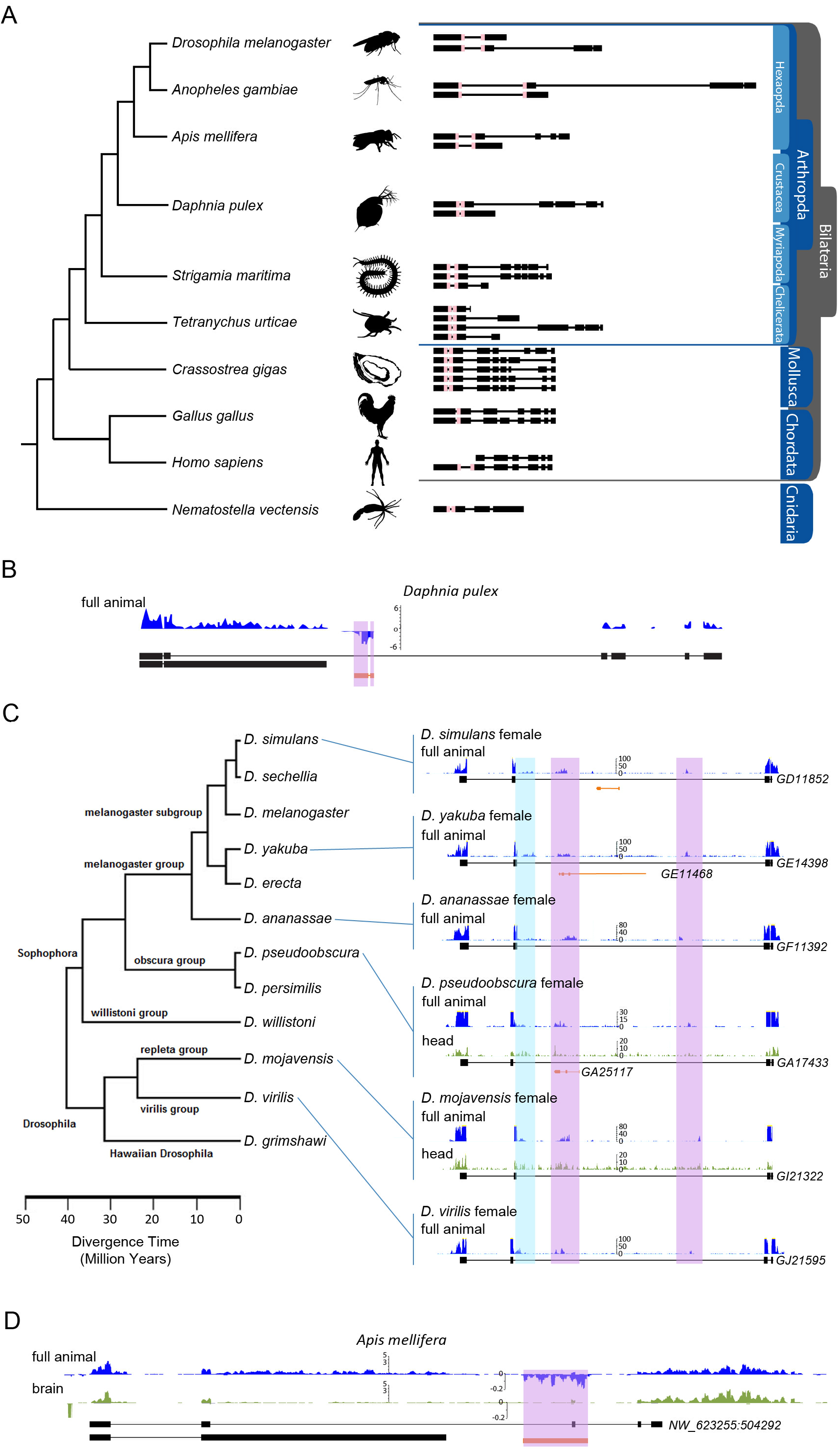
Evolutionary conservation of *bs* and *bsAS*. See also Supplementary Figure 4. **A,** Annotation of *bs* isoforms along metazoans. *bs* orthologous genes were manually annotated taking advantage of available RNA-Seq data in all species depicted. The pink box represents the MADS box, conserved along evolution. **B,** Expression of *bs* and *bsAS* in *Daphnia pulex*. An antisense transcript is detected within the long intron of *bs* gene. **C,** Expression of *bs* and *bsAS* in *Drosophila* species. Unstranded RNA-Seq tracks from modENCODE project are shown. When available, whole animals (in blue) and heads (in green) RNA-Seq samples have been represented. The long isoform of *bs* is annotated in all species. We have been able to identify reads likely corresponding to the short isoform of *bs* (highlighted in blue). *D. simulans*, *D. yakuba* and *D. pseudoobscura* also present annotated antisense genes embedded into the long intron of the coding gene (in orange). In all species, reads corresponding to the *bsAS* gene have been identified (in purple). **D,** Expression of *bs* and *bsAS* in *A. mellifera* whole animals (in blue) and brains (in green). As in fly, *bs* and *bsAS* are less expressed in brains compared to whole animals.

## Online Methods

### *Drosophila* strains

Fly strains used for this work are: wild-type (CantonS), *rn-GAL4/UAS-GFP*, *sal*^*E/Pv*^-*GAL4*, *bs-GAL4*, *bsAS*−/−, *UAS-bsAS*, *UAS-bsIsoB*, *UAS-bsLongIsos* and *w-; If/CyO; MKRS/TM6b*. Flies were grown in standard media at 25°C.

### *In situ* hybridization and immunohistochemistry

*In situ* hybridizations and immunostaining were carried out according to standard protocols. For *in situ* hybridization, *bsAS* DNA probe was synthesized by conventional PCR using a PCR DIG Labeling Mix (Roche). A biotinylated sense primer was used to bind the PCR product to streptavidin beads. Anti-sense probe was purified by denaturalization of the DNA attached to the beads. Primers used for probe synthesis are listed in Supplementary Table 4. Peroxidase conjugated anti-digoxigenin and Tyramide signal amplification (TSA, Life Technologies) were used for fluorescent *in situ* hybridization (FISH). DSRF immunostaining was performed with mouse anti-DSRF (Active Motif, 1:200). Fluorescently labeled secondary antibody was from Life Technologies. Discs were mounted in SlowFade (Life Technologies) supplemented with 1 μM DAPI (Life Technologies) to label nuclei. Wild type and *bsAS*−/− third instar larvae wing discs were analyzed with a SPE confocal microscope (Leica). For all *in situs* and immunostainings around 10 imaginal discs were analyzed. All experiments were performed twice.

### Chromatin immunoprecipitation

As starting material, 100 wt and *bsAS*−/− third instar larvae wing discs were used per experiment. Imaginal discs were manually dissected and pooled in 1 mL PBS-0.01 % TritonX100. Formaldehyde was added to a final concentration of 1% and tissues were fixed for 10 minutes at room temperature in a rotating wheel. Sonication was performed in a Diagenode Bioruptor for 15 minutes at high intensity with ON/OFF alternate pulses of 30 seconds. Sheared chromatin was aliquoted and flash frozen in liquid nitrogen. Chromatin immunoprecipitation assays were next performed as previously described(Perez-Lluch et al., 2011). For immunoprecipitation, 2 μg of anti-H3K4me3 (Abcam/ab8580) were used. Enrichments were analyzed by Real-Time PCR and primers are listed in Supplementary Table 4. Three independent biological replicates were performed per experiment.

### RNA isolation, library preparation and sequencing

RNA from imaginal discs was extracted with ZR-RNA MicroPrep Kit from Zymo Research following the manufacturer’s instructions. Sequencing libraries were prepared using TruSeq Stranded mRNA Library Preparation Kit from Illumina and following the manufacturer’s instructions. Sequencing was performed in a HiSeq sequencer from Illumina at the Ultrasequencing Unit of the Centre for Genomic Regulation (CRG, Barcelona, Spain). A minimum of 50 million paired-end 75 bp-long reads were obtained per replicate and two replicates were performed per each tissue.

### Retro-transcription and Real-Time PCR analyses

Retro-transcriptions and qPCRs were performed as described previously (Perez-Lluch et al., 2011). Primers used for Real-Time PCR are listed in Supplementary Table 4. Three independent biological replicates were performed per experiment.

### CRISPR-CAS9-induced deletion in flies

Guide RNAs around the *bsAS* TSS were designed using the CRISPR Target Finder from the flyCRISPR portal (http://flycrispr.molbio.wisc.edu/). Sequences of the gRNAs are listed in Supplementary Table 4. gRNAs were cloned into a *BbsI* digested pU6-*BbsI*-chiRNA vector(Gratz et al., 2013) following the protocol in flyCRISPR portal. A mixture of pU6-*BbsI*-gRNA1 and pU6-*BbsI*-gRNA2 was injected into embryos expressing CAS9 protein under the control of the *vasa* driver. Injected flies (F0) were crossed in groups of 4 males and females. F1 flies were allowed to mate and ley eggs for 10 days before screening. Sequential crosses were performed until the flies presenting the deletion were identified and isolated. Mutant flies were crossed with flies carrying a CyO balancer and homozygous mutants were isolated (Figure S2B). Flies were screened by conventional PCR. Briefly, genomic DNA was extracted from groups of 4 flies by smashing the animals in lysis buffer containing 0.5% NP40, 10 mM TrisHCl pH8.0, NaCl 150 mM, EDTA 2 mM and proteinase K 1 mg/mL. Genomic DNAs were incubated for 2 h at 50°C and centrifuged at top speed for 5 minutes to remove the remaining fly fragments. 2 μL of the extracts were used for each PCR. Primers used for the screening are listed in Supplementary Table 4.

### Transgenic constructs

For transgenesis, *bs* isoform B and long isoform (common to isoforms A and C) cDNAs were amplified by rtPCR (primers used for the amplifications are listed in Supplementary Table 4) and inserted by Gibson cloning into a pUAST-AttB vector digested with *EcoRI*. Transgenes were directed into phi31 fly strain J36, in the chromosome III.

### Chromosome Conformation Capture (3C)

Chromosome conformation capture protocol was adapted from previous reports(Naumova et al., 2012). For each experiment a 3C library and a genomic library were performed. For genomic libraries, genomic DNA from 5 third instar larvae was extracted by smashing the animals in lysis buffer containing 0.5% NP40, 10 mM TrisHCl pH8.0, NaCL 150 mM, EDTA 2 mM and proteinase K 1 mg/mL. Genomic DNA was incubated for 2 h at 50°C, centrifuged at top speed for 5 minutes to remove the remaining larval fragments, purified by performing two rounds of phenol:chloroform extraction and precipitated with ethanol. For 3C libraries, 150 wing imaginal discs were manually dissected and pooled in 1 mL PBS-0.01 % TritonX100. Formaldehyde was added to a final concentration of 1% and tissues were fixed for 10 minutes at room temperature in a rotating wheel. Discs were washed in 1 mL of PBS-0.01 % TritonX100-0.125 M glycine for 5 minutes at room temperature and finally resuspended in 500 μL of lysis buffer (10 mM Tris pH 8, 10 mM NaCl, 0.2% NP40 1× Complete Protease Inhibitor Cocktail) and flesh frozen in liquid nitrogen. To perform the libraries, 5 μg of genomic DNA and the fixed 150 wing discs were equilibrated in 164 μL of digestion mix (20 μl *HhaI* 10× buffer, 5 μl 10% SDS and 139 μl H_2_O) for 1 h at 37° in a ThermoMixer. 32 μl of 10% TritonX100 were added to each library and incubated for 1 h more at 37°C. Digestion was performed by adding 2 mL of *HhaI* restriction enzyme (New England Biolabs) and incubating for 2 h at 37°C. After first digestion, 2 μl of restriction enzyme were added and samples were incubated over night at 37°C. Digestions were inactivated for 20 minutes at 65°C and cooled on ice. To perform the ligation, 165 μL of digestion mix (2× T4 ligation buffer + 4 μl T4 ligase) were added to the libraries. Ligations were incubated for 4 h at 18°C. Libraries were de-crosslinked by adding 2 μl of proteinase K 20 mg/mL and incubating over night at 65°C. Remaining RNAs in the samples were removed by RNase treatment for 30 minutes at 37°C. Libraries were finally purified by phenol:chloroform extraction and precipitated in ethanol. Quantification of interactions was performed by Real-Time PCR, comparing the interaction observed in the regions of interest to a known interacting region (positive control). Primers used for Real-Time PCRs are listed in Supplementary Table 4. At least, three independent biological replicates were performed per experiment.

### RNA-Seq processing

Data was processed using grape-nf (available at https://github.com/guigolab/grape-nf). RNA-Seq reads were aligned to the fly genome assembly dm6 (dos Santos et al., 2015) using STAR 2.4.0 software (Dobin et al., 2013) with up to 4 mismatches per paired alignment using the FlyBase genome annotation r6.05 (Attrill et al., 2016). Only alignments for reads mapping to ten or fewer loci were reported. Gene and transcripts TPMs were quantified using RSEM (Li and Dewey, 2011). Tracks were visualized as a Track Hub at the UCSC Genome Browser(Raney et al., 2014).

### NATs in the fly genome and in larval tissues

Antisense lncRNAs were inferred from their overlap and orientation relative to protein coding genes. A minimum overlap of 1 exonic nucleotide was required to classify the pairs into the following configurations: (i) 5’ head-to-head: first exon overlap; (ii) 3’ tail-to-tail: last exon overlap; (iii) internal: lncRNA is within the protein coding gene; (iv) external: protein coding gene is within the lncRNA; and (iv) complex: overlap of more than 2 genes.

Pairs of ncRNAs antisense to mRNAs with more than 1 isoform and expressed at least 0.5 TPMs in at least both replicates of any L3 sample were selected for further analyses. Ratio of exon coverage over the gene coverage was computed using bwtool summary (Pohl and Beato, 2014). Candidates pairs were ranked based on the Pearson's coefficient of correlation of exon ratio and antisense expression and on the standard deviation of exon ratios. Exons showing a standard deviation higher than 0.1 and an absolute correlation higher than 0.7 were selected.

### Differential gene expression analysis of mutant versus wild type samples

Pairwise differential gene expression (DEG) analysis between wild type and mutant samples was performed using EdgeR (Robinson et al., 2010). Only genes expressed at least 5 TPM in at least two samples were selected for this analysis. DEG selected showed log_2_ fold change > 2 and FDR < 0.01. Biological Processes from Gene Ontology database enriched in upregulated and downregulated datasets were assessed by using GOstats(Falcon and Gentleman, 2007) and visualized with ReviGO(Supek et al., 2011), with a p value cutoff of 0.001.

### Promoter and ChIP-Seq analysis

To search for transcription factor binding sites (TFBSs) in *bs* and *bsAS* TSSs we used the Promo 3.0 tool(Farre et al., 2003). We selected TFBSs with a matrix dissimilarity rate less or equal than 5% were present both in *bs* TSS1 and *bsAS* TSS but not in *bs* TSS2. Available ChIP-Seq data of GAF(Oh et al., 2013), RNA PolII(Schertel et al., 2015), Pc and Ph(Loubiere et al., 2016) were aligned to the fly genome (dm6) using GEM-Mapper(Marco-Sola et al., 2012) with up to 2 mismatches per read using the dm6 genome assembly(Attrill et al., 2016; dos Santos et al., 2015). ChIP-Seq data from modENCODE TFs were visualized at UCSC Genome Browser from the ENCODE Portal(Davis et al., 2018).

### Evolutionary conservation

To track down the evolutionary conservation of *bs* and *bsAS*, we first analyzed RNA-Seq data of whole body and head of female *Drosophila* species from modENCODE (GSE44612(Chen et al., 2014)). Next we investigated the expression of *bs* and *bsAS* in stranded RNA-Seq data of *Apis mellifera* (whole body: GSE83437(Mao et al., 2017) and head: GSE87001(Shpigler et al., 2017); UCSC apiMel2 assembly; modelRefGene genome annotation available at UCSC), in stranded RNA-Seq data of *Anoplophora glabripennis* (PRJNA274806;) in unstranded RNA-Seq data of *Folsomia candida* (PRJNA239929; RefSeq assembly accession: GCF_002217175.1; NCBI *Folsomia candida* Annotation Release 100) and in stranded RNA-Seq of *Daphnia pulex* (GSE103939; assembly GCA_000187875.1; annotation ENSEMBL release-26). Finally, we manually annotated *bs* isoforms across metazoans taking advantage of available RNA-Seq data and relying on split reads to properly annotate the exon junctions, which were visualized with ggsashimi(Garrido-Martin et al., 2018): *Anopheles gambiae* (GSE55453(Vannini et al., 2014); GSE59773(Gomez-Diaz et al., 2014)), *Apis mellifera* (GSE52289(Cameron et al., 2013); GSE65659(Galbraith et al., 2015); SRP068487), *Daphnia pulex* (DRP002580), *Strigamia maritima* (SRP041623), *Tetranychus urticae* (GSE31527(Grbic et al., 2011; Van Leeuwen et al., 2012)), *Crassostrea gigas* (GSE31012(Zhang et al., 2012); SRP058882), *Gallus gallus* (GSE41338(Barbosa-Morais et al., 2012)), and *Nematostella vectensis* (GSE46488(Schwaiger et al., 2014)). *Homo sapiens bs* annotation was directly extracted from GENCODE v19(Frankish et al., 2018). Data was processed using grape-nf (available at https://github.com/guigolab/grape-nf) with the same parameters as our RNA-Seq data.

## Author contribution

S.P.L., A.B. and R.G. conceived and designed the study and the experiments. S.P.L., M.R.R., A.A., C.A. and L.B. performed the experiments. S.P.L., A.B. and C.K. performed the computational analyses with the assistance of E.P. S.P.L., A.B., C.K. and R.G. wrote the manuscript with contributions from all authors.

## Acknowledgments

We thank Bruna Correa, Ramil Nurdinov, Sebastian Ullrich and Beatrice Borsari for helpful discussions about the data and the manuscript and Romina Garrido for administrative assistance. We also thank the Ultrasequencing Unit and the Advanced Light Microscopy Unit of the CRG (Barcelona, Spain), for sample processing. We also thank Montserrat Corominas and Florenci Serras, from the University of Barcelona, for the anti-DSRF antibody. This work was performed under the financial support of the European Community under the FP7 program (ERC-2011-AdG-294653-RNA-MAPS), from the Spanish Ministry of Economy and Competitiveness (MEC) under grant number BIO2011-26205 and the Centro de Excelencia Severo Ochoa, from the CERCA Programme (Generalitat de Catalunya). This research reflects only the authors’ views and the Community is not liable for any use that may be made of the information contained therein.

## Data access

RNA-Seq raw data and processed files were deposited in the ArrayExpress database at EMBL-EBI (www.ebi.ac.uk/arrayexpress) under accession number E-MTAB-7653.

## Supplementary Figures

**Supplementary Figure 1.**
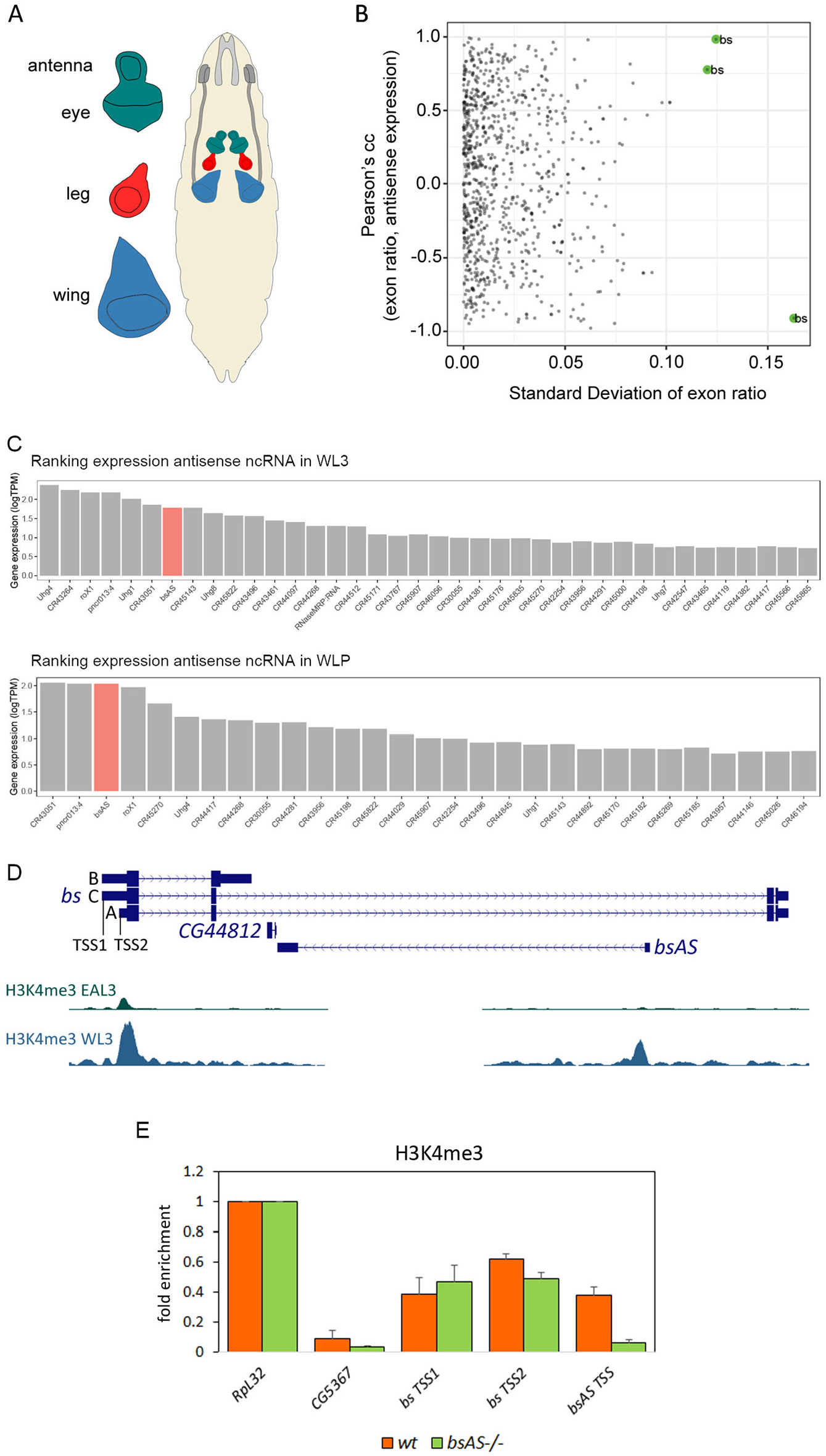
*bsAS* is mainly expressed in *Drosophila melanogaster* wing tissues. Related to figures 1 and 2. **A,** Third instar larvae tissues used for RNA-Seq experiments. **B,** Relationship between the correlation of exon ratio and antisense expression and the standard deviation (st. dev.) of sense gene exons ratio. Of the 145 SA pairs with both genes expressed in at least one tissue in L3, 104 pairs (72%, 102 genes) involve a protein coding gene with multiple isoforms. We computed the correlation between the expression of these NATs and the inclusion of the 964 exons of the sense protein coding genes. Because of the very small number of independent data points, we have little power to detect significant correlations. Still, we plotted this correlation against the standard deviation of the exon inclusion values, to focus specifically in the exons that changed the most (see Methods). Among them, the case of *blistered* -*bs*- is the strongest, as it indeed shows strong positive and negative correlation between the expression of the NAT antisense to *bs* and the inclusion of three highly variable *bs* exons. **C,** Ranking of expression of antisense lncRNAs in L3 (upper panel) and LP (lower panel) wings. *bsAS* is highlighted in red. **D,** Histone methylation marks (Perez-Lluch et al., 2015) at *bs* locus. There is a strong peak of H3K4me3 at *bs* TSS and *bsAS* TSS in WL3, whereas only a small peak at *bs* TSS is observed in EAL3. **E,** H3K4me3 ChIP-qPCR in WL3. No differences are observed between *bs* TSS1 and TSS2 H3K4me3 marking. The deletion of *bsAS* TSS induces a dramatic reduction of H3K4me3 marking in *bsAS* TSS. TSS1 and TSS2 of *bs* do not show differences between *bsAS*−/− and wt. *RpL32* and *CG5367* correspond to positive and negative controls of the ChIP, respectively.

**Supplementary Figure 2.**
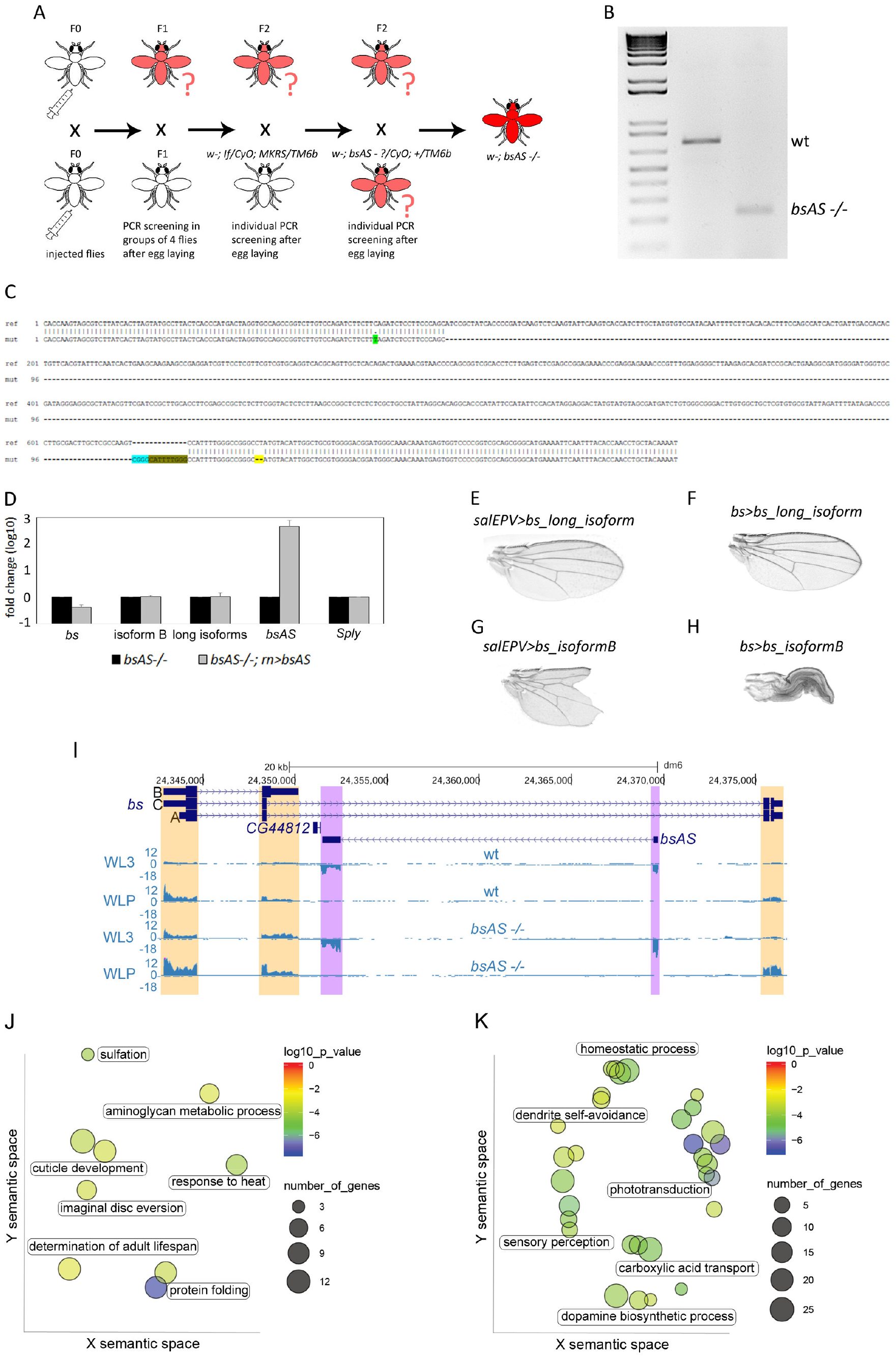
*bsAS* depletion induces strong phenotype in wings and overexpression of neural genes. Related to figure 2. **A,** *Drosophila* embryos expressing CAS9 nuclease in the germinal cells under the control of *vasa* driver, were injected with a mix of two plasmids expressing gRNAs against the TSS of *bsAS*. The screening was performed retroactively, by allowing putatively mutant flies to lie eggs before screening. Once the mutation was isolated, genetic crosses were performed to obtain the homozygous mutant flies. **B,** PCR screening of *bsAS* deletion. A band of 800 bp was amplified in wt flies, whereas a band of 250 bp was obtained from homozygous *bsAS* mutant flies. **C,** Alignment of wt and *bsAS*−/− genomic regions. A Single Nucleotide Polymorphism was detected in the *bsAS* TSS sequence (in green). The 5’ region of the deletion has been cleanly repaired, but in the 3’ region, several insertions/deletions have occurred (the insertion of 4 nucleotides –in blue-, the duplication of 9 nucleotides –in green-; and the deletion of 2 nucleotides –in yellow-). **E,** Overexpression of *bsAS* under the control of *sal*^*EP/V*^ driver (specific driver that expresses GAL4 restricted to the the *salm* domain within the wing pouch), checked by *in situ* hybridization. Upper panels, *bsAS* (in red) is overexpressed in the central part of the wing, indicating that the designed *bsAS* probe specifically hybridizes with *bsAS*. Lower panels, control sense probe (also in red) gives no signal in the wing pouch. **E-H,** Overexpression of *bs* isoforms in wings. The overexpression of long isoforms of *bs* under the control of *sal*^*E/Pv*^ (**E**) and *bs* (**F**) drivers do not show any evident defect in wing morphology and vein patterning. **G,** The overexpression of the *bs* isoform B under the s*al*^*E/Pv*^ driver induces the appearance of notches in the wing margin. **H,** The overexpression of the isoform B of *bs* under the control of *bs* driver induces high lethality, likely due to the expression of this driver in the Central Nervous System. The few *bs>bs_isoformB* escapers obtained present almost unfolded wings, with no obvious vein/intervein patterning. I, Expression derived from RNA-experiments of *bs* locus in wt and *bsAS* −/− wings at L3 (upper panels) and LP (lower panels) –note that wt tracks coincide with tracks in Figure 1A–. Differences in expression between wt and mutant tissues are depicted in purple for the lncRNA and in green and blue for the TSS and the TTS of the coding gene, respectively. The depletion of *bsAS* expression after *bsAS* TSS deletion is confirmed in the RNA-Seq experiments, as well as the consequent overexpression of the *bs* isoform C. J, Gene Ontology Term Enrichment analysis of downregulated genes in *bsAS*−/− WLPs. Few GO categories were enriched among downregulated genes in *bsAS*−/− LP wings, among them, cuticle development and imaginal disc eversion. **K,** Gene Ontology Term Enrichment analysis of upregulated genes in *bsAS*−/− WLPs. Upregulated genes are enriched in categories mainly related to neural fates, such as phototransduction, sensory perception and dopamine biosynthesis. In **J** and **K,** each spot represents an enriched category and labels are representative of clusters of categories; color of the spots represents the p value of enrichment; size of the spots represents number of genes belonging to each category. Axes represent semantic space from ReviGO (Supek et al., 2011).

**Supplementary Figure 3.**
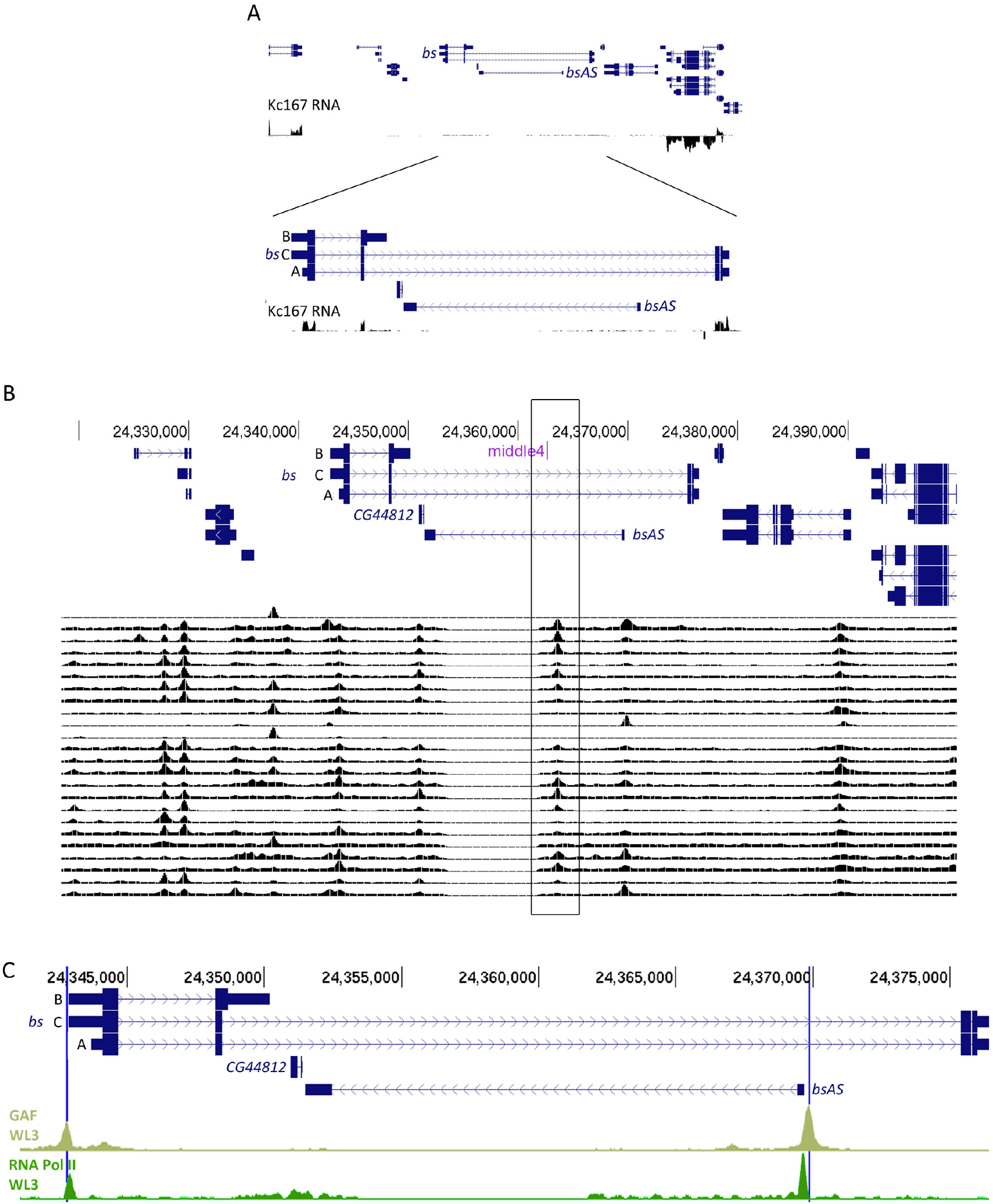
Interactions along the *bs* locus. Related to figure 3. **A,** RNA-Seq of Kc167 cells, from modENCODE (Duff et al., 2015). *bsAS* is not expressed in these cells. *bs* expression is very low and isoform A seems to be main one expressed. **B,** Profile of DNA binding of 24 transcription factors during embryogenesis from modENCODE project (Washington et al., 2011). Middle4 is very close to a hot spot for transcription factor binding. **C,** Profile of DNA binding of GAF (Oh et al., 2013) (light green) and RNA Pol II (Schertel et al., 2015) (dark green) in third instar larvae wings. GAF and RNA Pol II peaks coincide with the expression of *bs* TSS1 and *bsAS* in wings. Blue vertical lines represent GAF binding sites at *bs* TSS1 and *bsAS* TSS.

**Supplementary Figure 4.**
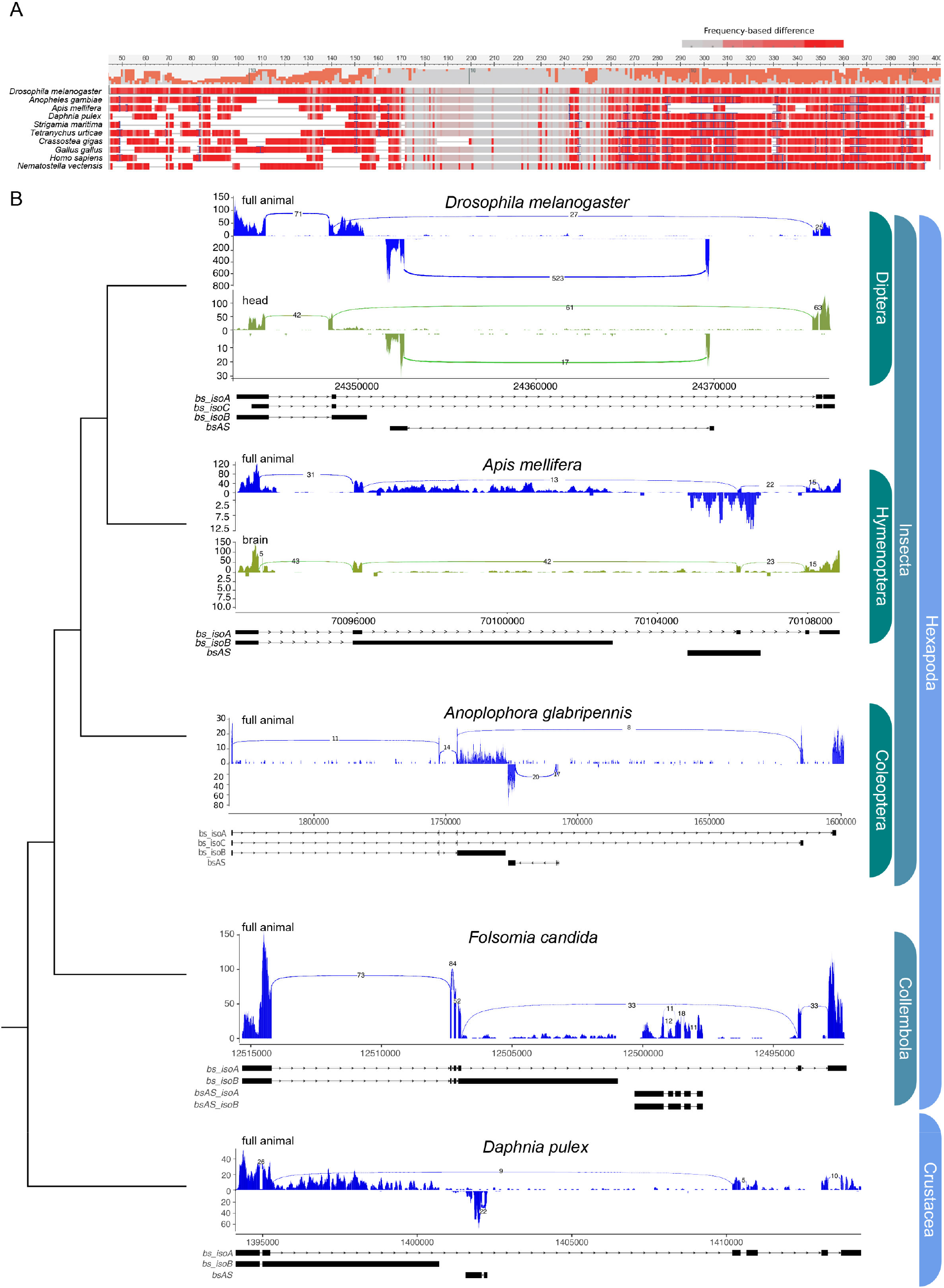
Evolutionary conservation of *bs* and *bsAS*. Related to figure 4. **A,** Multiple sequence alignment of the long *bs* isoform using MAFFT and visualization of the frequency-based difference using NCBI MSA Viewer. High sequence conservation is observed between 160-270bp where MADS-box is located. Sequence conservation drops rapidly outside this region. **B,** Read-depth along *bs* locus. Organisms are sorted by the tree of life. Number of split reads are highlighted in the exon junctions generated using ggsashimi (Garrido-Martin et al., 2018). Stranded RNA-Seq is shown in two separated strands, where the negative strand is negated and shown below the positive strand. When available, whole animals (in blue) and heads (in green) RNA-Seq samples have been represented. The long isoform of *bs* is annotated in all species. Expression of both long and short isoforms of *bs* is supported by read coverage and by split alignments in exon junctions in represented species. Antisense expression is supported by stranded read coverage and by split alignments in the exon boundaries in the Diptera *Drosophila melanogaster*, in the Coleoptera *Anoplophora glabripennis* and in the Crustacea *Daphnia pulex*. Stranded RNA-Seq of *Apis mellifera* presents more antisense signal in the whole body compared to the head sample, however there are no split reads supporting exon junctions. The wingless basal Hexapoda, *Folsomia candida*, presents a clear set of split reads supporting the junctions of *bsAS* and no split reads shared between *bs* and *bsAS*, however the data is unstranded. Overall, Hexapoda and Crustacea show expression of both long-short *bs* isoforms as well as the antisense expression on the lncRNA *bsAS*.

## Supplementary Tables

**Supplementary Table 1.** Summary statistics of RNA-Seq samples. Related to figures 1 and 2. All samples generated along the manuscript are summarized here.

| Sample | Total # reads | # Mapped reads | Proportion mapped reads | # Uniquely mapped reads | Proportion uniquely mapped reads |
| --- | --- | --- | --- | --- | --- |
| wtEAL3.1 | 92624048 | 91486808 | 98.7722 | 88894992 | 97.16701 |
| wtEAL3.2 | 77533626 | 75828528 | 97.80083 | 73725748 | 97.22693 |
| wtLL3.1 | 121021466 | 107715318 | 89.00513 | 105383724 | 97.83541 |
| wtLL3.2 | 109413972 | 102201120 | 93.40774 | 97882412 | 95.7743 |
| wtWL3.1 | 81030122 | 79816358 | 98.50208 | 78079790 | 97.8243 |
| wtWL3.2 | 83521572 | 82426862 | 98.68931 | 78321148 | 95.01896 |
| wtELP.1 | 134304082 | 129152790 | 96.16446 | 126591202 | 98.01662 |
| wtELP.2 | 132043336 | 121297912 | 91.8622 | 118183372 | 97.43232 |
| wtWLP.1 | 108100128 | 105568376 | 97.65796 | 102489310 | 97.08334 |
| wtWLP.2 | 144086210 | 136518300 | 94.74765 | 133541172 | 97.81925 |
| bsEAL3.1 | 147819126 | 136876976 | 92.59761 | 134435818 | 98.21653 |
| bsEAL3.2 | 139808904 | 106272586 | 76.01275 | 85335706 | 80.29889 |
| bsWL3.1 | 184430738 | 164711012 | 89.30779 | 160623490 | 97.51837 |
| bsWL3.2 | 139062704 | 136397368 | 98.08336 | 133643534 | 97.98102 |
| bsELP.1 | 210915056 | 201795336 | 95.67612 | 198332716 | 98.28409 |
| bsELP.2 | 206900922 | 197297590 | 95.35849 | 193799048 | 98.22677 |
| bsWLP.1 | 236383298 | 224236494 | 94.86139 | 220943980 | 98.53168 |
| bsWLP.2 | 223774820 | 211371452 | 94.45721 | 207968748 | 98.39018 |

**Supplementary Table 2.** Gene and transcript expression of *bs* and *bsAS* in wt and *bsAS*−/− tissues. Related to figures 1 and 2. Genes are highlighted in grey. Expression values are represented in TPMs.

|  |  | wt |  |  |  | *bsAS*<sup>-/-</sup> |  |  |  |
| --- | --- | --- | --- | --- | --- | --- | --- | --- | --- |
| gene | Gene/transcript | WL3 | WLP | EAL3 | ELP | WL3 | WLP | EAL3 | ELP |
| *blistered* | FBgn0004101 | 25.545 | 48.175 | 7.81 | 3.87 | 72.68 | 120.79 | 9.885 | 9.405 |
|  | FBtr0072271 (A) | 1.135 | 4.035 | 7.275 | 2.985 | 12.815 | 16.215 | 9.625 | 4.755 |
|  | FBtr0290089 (B) | 18.89 | 42.21 | 0.535 | 0.645 | 16.13 | 34.965 | 0.06 | 0.845 |
|  | FBtr0343314 (C) | 5.52 | 1.93 | 0 | 0.24 | 43.735 | 69.61 | 0.2 | 3.805 |
| *bsAS* | FBgn0266046 | 60.185 | 108.79 | 2.145 | 1.585 | 0.03 | 0.075 | 0 | 0.02 |
|  | FBtr0343311 | 60.185 | 108.79 | 2.145 | 1.585 | 0.03 | 0.075 | 0 | 0.02 |

**Supplementary Table 3. DEG between wt and *bsAS*−/− in L3 and LP wings.** Related to figure 2. Expression values are represented in TPMs. L3 DEG are represented in sheet 1 and LP DEG are represented in sheet 2.

**Supplementary Table 4. List of primers used to perform the study.** Related to figures 1, 2 and 3. Name of the primer, sequence and experiment in which it has been used are depicted.

